# Differential geometry methods for constructing manifold-targeted recurrent neural networks

**DOI:** 10.1101/2021.10.07.463479

**Authors:** Federico Claudi, Tiago Branco

**Affiliations:** Sainsbury Wellcome Centre for Neural Circuits and Behaviour, UCL, London, UK

## Abstract

Neural computations can be framed as dynamical processes, whereby the structure of the dynamics within a neural network are a direct reflection of the computations that the network performs. A key step in generating mechanistic interpretations within this *computation through dynamics* framework is to establish the link between network connectivity, dynamics and computation. This link is only partly understood. Recent work has focused on producing algorithms for engineering artificial recurrent neural networks (RNN) with dynamics targeted to a specific goal manifold. Some of these algorithms only require a set of vectors tangent to the target manifold to be computed, and thus provide a general method that can be applied to a diverse set of problems. Nevertheless, computing such vectors for an arbitrary manifold in a high dimensional state space remains highly challenging, which in practice limits the applicability of this approach. Here we demonstrate how topology and differential geometry can be leveraged to simplify this task, by first computing tangent vectors on a low-dimensional topological manifold and then embedding these in state space. The simplicity of this procedure greatly facilitates the creation of manifold-targeted RNNs, as well as the process of designing task-solving on-manifold dynamics. This new method should enable the application of network engineering-based approaches to a wide set of problems in neuroscience and machine learning. Furthermore, our description of how fundamental concepts from differential geometry can be mapped onto different aspects of neural dynamics is a further demonstration of how the language of differential geometry can enrich the conceptual framework for describing neural dynamics and computation.

## Introduction

Networks of neurons can be viewed as dynamical systems in which the joint activity of all units is a state that represents the information stored in the network, and its dynamics represent computations **(Vyas et al., 2020; Sussillo, 2014)**. Under this *computation through dynamics* perspective, understanding neuronal computation requires describing the dynamics of neural networks and how these are determined by their connectivity structure **(Schaeffer et al., 2020; Finkelstein et al., 2021**). Neural dynamics are often conceptualized as trajectories in an *n*-dimensional vector space - the state space - in which the distance along each dimension represents the firing rate of individual neurons. Recent work suggests that in both biological and artificial neural networks, such neural trajectories are often confined to a lower dimensional subspace of state space **(Gao and Ganguli, 2015; Gao et al., 2017; Russo et al., 2018; Gallego et al., 2017; Maheswaranathan et al., 2019)**, and occasionally, to a neural manifold with additional topological structure **(Kim et al., 2017; Gardner et al., 2021; Chaudhuri et al., 2019)**. It has been further hypothesized that the geometry and topology of these low dimensional neural manifolds is linked in a fundamental way to the computations carried out by the network **(Maheswaranathan et al., 2019; Gao et al., 2017; Jazayeri and Ostojic, 2021; Darshan and Rivkind, 2021; Chung and Abbott, 2021; Pollock and Jazayeri, 2020)**. This view therefore emphasizes that understanding how network connectivity gives rise to structured neural dynamics is a key goal towards explaining neural computations.

Achieving this goal will likely require the measurement of activity and connectivity of large number of neurons spanning multiple brain regions in individual animals. While recent technological developments have enabled simultaneous activity recordings from hundreds of neurons and detailed reconstructions of network connectivity **(Steinmetz et al., 2021; Stringer et al., 2019; Winnubst et al., 2019; Osten and Margrie, 2013)**, a complete description of a network’s structure and activity in behaving animals remains largely beyond the reach of experimental neuroscience. On the other hand, artificial Recurrent Neural Networks (RNNs) can be trained to solve a variety of tasks similar to those employed in experimental neuroscience, and their connectivity structure and dynamics are perfectly known. This makes them an ideal testing ground for developing theoretical and analytical tools to investigate the links between connectivity, dynamics and computation **(Sussillo, 2014; Sussillo and Barak, 2013; Mastrogiuseppe and Ostojic, 2018)**. The majority of work employing RNNs to address these issues uses tools from dynamical systems theory to reverse-engineer the neural dynamics of networks trained to perform a task **(Sussillo and Barak, 2013; Schaeffer et al., 2020)**. This optimization-based approach allows RNNs to discover how to best structure their dynamics to carry out a certain computation. An alternative approach is to develop general algorithms for directly constructing networks with dynamics that are targeted to a pre-selected manifold, in an effort to produce a deeper understanding of how connectivity determines dynamics **(Mastrogiuseppe and Ostojic, 2018; Beiran et al., 2020; Eliasmith and Anderson, 2003; Pollock and Jazayeri, 2020; Biswas and Fitzgerald, 2020; Darshan and Rivkind, 2021)**. A particularly promising approach in this direction is the Embedding Manifolds with Population-level Jacobians (EMPJ) algorithm **(Pollock and Jazayeri, 2020)**, which enables the creation of RNNs with dynamics confined to a target manifold in state space. Briefly, EMPJ uses vectors tangent to the target manifold to build a system of equations that yields the connectivity matrix for a network with dynamics that lay on the target manifold. Tangent vectors play a dual role in EMPJ: by being tangent to the target manifold they confine the RNN dynamics to it, and their orientation and magnitude dictate the on-manifold dynamics. EMPJ is therefore an elegant and parsimonious algorithm capable of producing RNNs targeted to different manifolds and displaying rich on-manifold dynamics.

In practice, however, computing the tangent vectors may be far from trivial, which effectively limits the range of problems that methods like EMPJ can be applied to. For example, a target manifold of given topology (e.g., the plane) can be embedded in state space in infinite ways. This makes computing tangent vectors a challenging task since the position and orientation of tangent vectors depend on the precise geometry of the target manifold (**Figure 1**A). The problem is made even harder by the necessity to precisely orient the tangent vectors to produce the desired on-manifold dynamics (e.g., to create attractor states; Figure **Figure 1**B, bottom). Here we aim to address this challenge by describing a general approach that simplifies the task of computing tangent vectors on target manifolds and specifying the required on-manifold dynamics (Figure **Figure 1**B). We employ concepts from topology and differential geometry to show how tangent vectors can be computed on topological manifolds prior to their embedding in state space (Figure **Figure 1**A, left), thus removing the need for recomputing the tangent vectors for different choices of manifold embeddings. This approach can in principle be combined with local geometry-based algorithms such as EMPJ, thus widening the class of problems that they can be readily applied to. We believe that this work adds to the view that ideas from differential geometry map naturally onto neural dynamics concepts and are therefore a powerful framework for understanding the link between connectivity, dynamics and computation.

**Figure 1.**
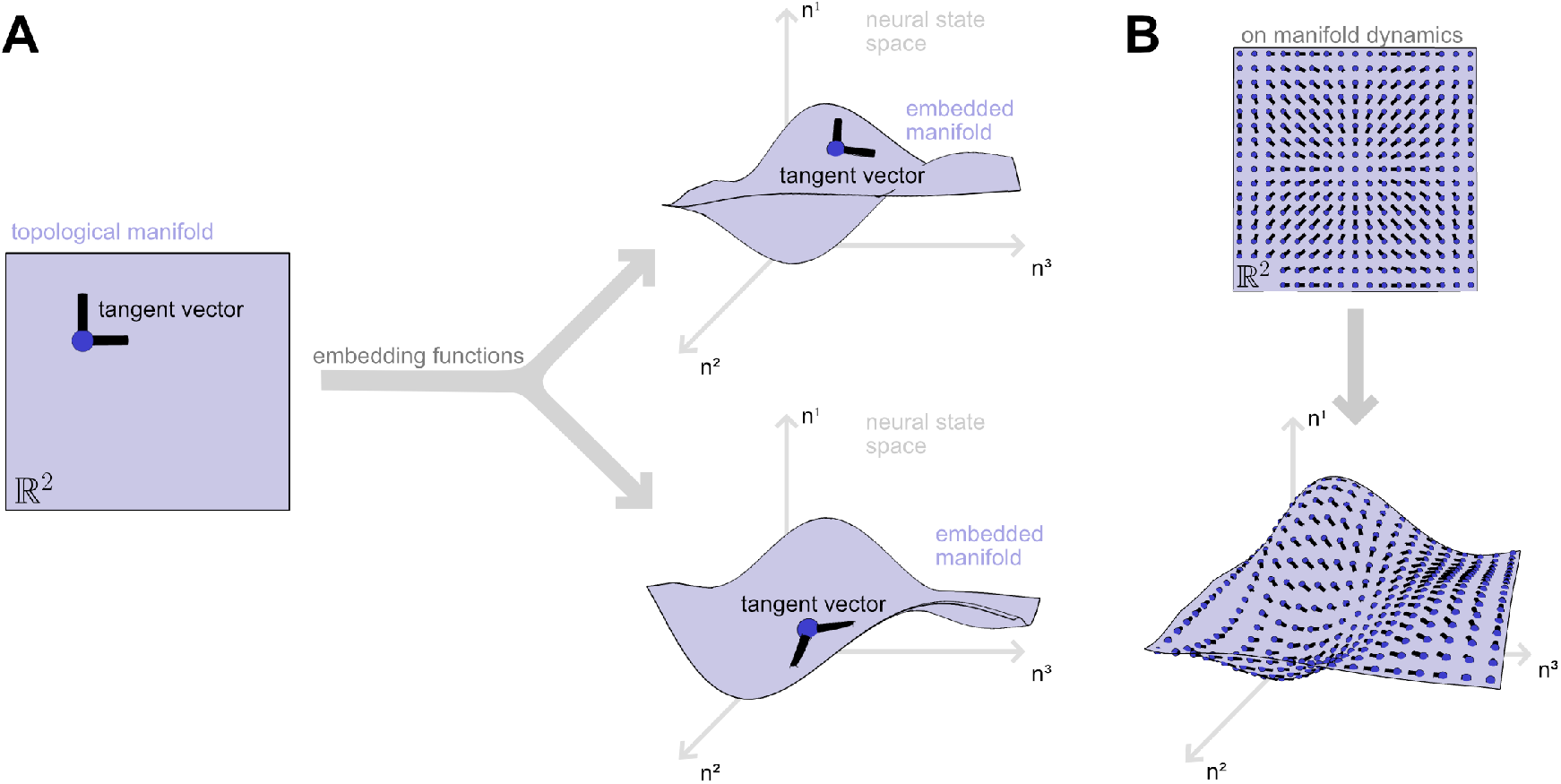
**A**. Left, the topological manifold ℝ^2^ with a schematic representation of tangent vectors at a point. Right, two different embeddings of ℝ^2^ in ℝ^3^ with the corresponding tangent vectors. **B**. On manifold dynamics. Top, schematic representation of a tangent vectors field on the topological manifold ℝ^2^. Bottom, the corresponding tangent vector field for an embedding of ℝ^2^ in ℝ^3^.

## Computing tangent vectors

In this section we demonstrate the application of concepts from differential geometry to the task of computing tangent vectors for creating manifold targeted RNNs. We leave the precise definitions of key objects, highlighted in bold, to a Mathematical Appendix section, and instead focus on providing an intuitive understanding of the method.

### Tangent vectors on the topological manifold

A **topological manifold** ℳ is a **topological space** locally homeomorphic to Euclidean space ℝ^*d*^(e.g.: the plane and the sphere are locally similar to ℝ^2^ at every point). In neuroscience, neural manifolds are often conceptualized as being situated or embedded in a higher dimensional vector space ℝ^*n*^, the neural state space. A topological manifold, however, does not necessarily exist within a larger space. Indeed, computations are often more easily carried out on the manifold prior to its embedding, like in the case of tangent vectors. For clarity, here we will refer to a manifold embedded in state space as an embedded manifold and as a topological manifold otherwise, although technically both are topological manifolds. A topological manifold is simply defined by a set and a **topology**. For example, the line ℝ^1^ manifold can be represented by the set *M* = [0, 1] endowed with the **standard topology**. We leave the definition of each of the manifolds used in this work to the Methods section.

For an *m*-dimensional manifold ℳ embedded in an *n*-dimensional space (with *n > m*), **tangent vectors** are *n*-dimensional vectors tangent to ℳ at a point. This view of tangent vectors, however, depends on the manifold being embedded in a larger vector space, and thus cannot be used as a general definition of a tangent vector on a topological manifold. Instead, tangent vectors at a point *p ∈ ℳ* are defined as equivalence classes of parametrized curves *γ* : ℝ ⊃ *I* → ℳ such that two curves are equivalent if they share the same directional derivative at *p*. This more abstract definition of tangent vectors is equivalent to the traditional view of tangent vectors for embedded manifolds (see below) but it enables us to compute tangent vectors on the topological manifold directly.

Any tangent vector *v*_*p*_ belongs to a tangent vector space at *p* : *T*_*p*_ *M* and can thus be defined as a linear combination of the basis vectors of *T*_*p*_*M*. A fundamental theorem in differential topology establishes that *dim*(*T*_*p*_ *M*) = *dim*(ℳ) such that *d*-many basis vectors need to be defined. The definition of the basis of *T*_*p*_ *M* relies on the concept of **chart** of a topological manifold: a chart establishes a local coordinates system by mapping an open neighborhood *U* ⊂ *M* to a subset of ℝ^*d*^, using a bijective map *x* (**Figure 2**A). For a point *p* it’s then possible to determine its position *x*(*p*) in the local coordinates system defined by a chart. In the same coordinates system it is possible to define *d*-many parametrized functions *f*_*i*_ going through *x*(*p*) and parallel to one axis of the local coordinates system. These can then be projected onto the manifold as *x*^−1^ ◦ *f*_*i*_ giving a set of parametrized functions on the manifold with different directional derivatives which can act as representative functions for basis vectors of *T*_*p*_*M* (**Figure 2**A). Let *e*_*i*_ = [*x*^−1^ ◦ *f*_*i*_] be a basis vector, then any tangent vector *v*_*p*_ can be expressed as a linear combination of the basis vectors: 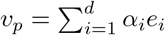 where the *α*_*i*_ are scalar factors.

**Figure 2.**
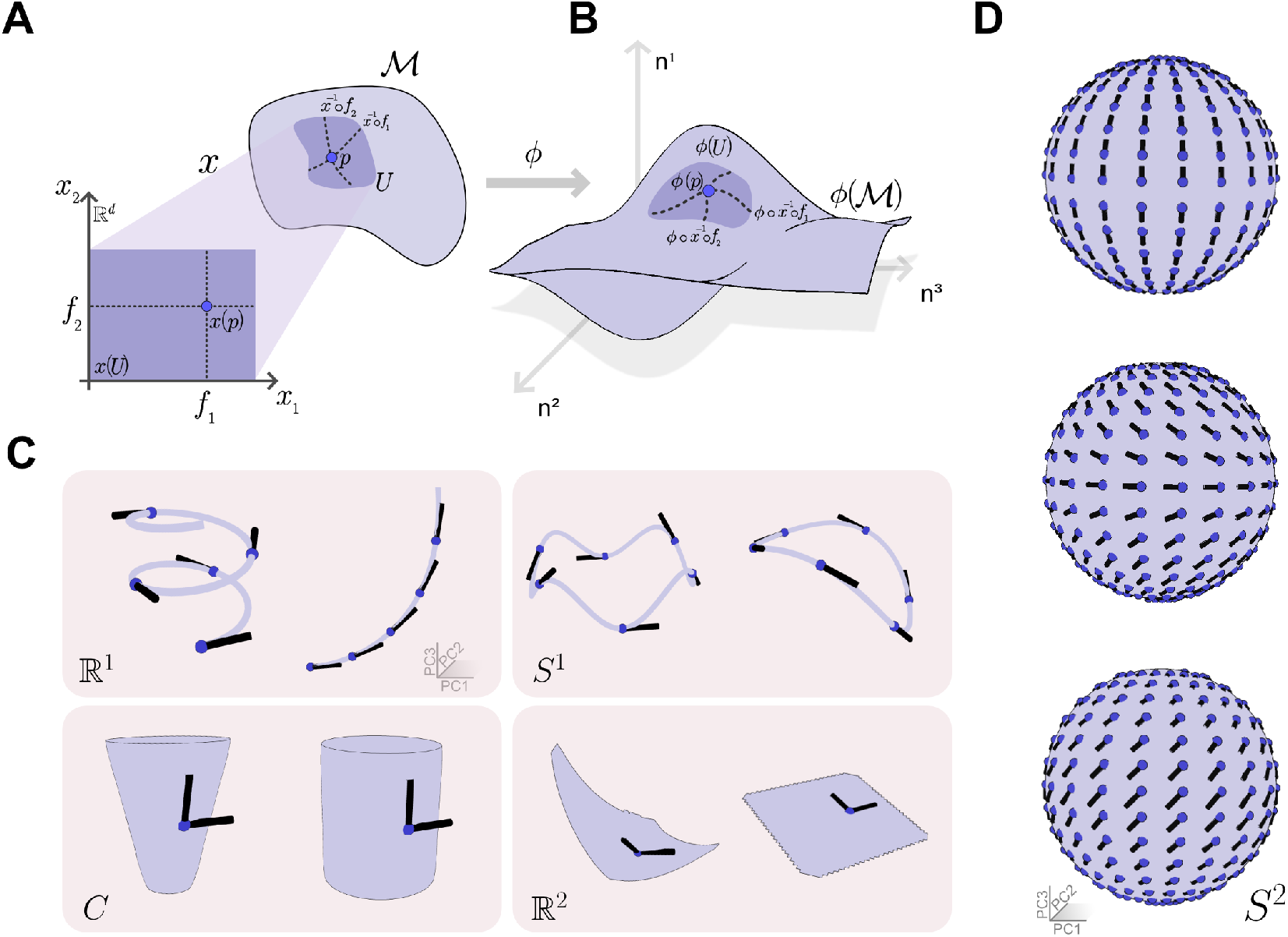
**A**. Graphic representation of a two dimensional topological manifold ℳ and a chart (*x, U*) containing a point *p*. The basis functions *f*_*i*_ in the chart’s local coordinates system are also shown. **B**. Graphic representation of an embedding of ℳ with the basis functions *ϕ x*^−1^ ◦ *f*_*i*_ shown. **C**. Tangent vectors (black) at selected points (blue dots) on one and two dimensional manifolds (light blue lines and surfaces). **D**. Tangent vector fields on the sphere. The tangent vectors produced by three different tangent vector fields for the sphere manifold are shown. For panels C and D, the manifolds were embedded in ℝ^64^ and visualized in three dimensions using the first three principal components of a PCA model, see Methods for details.

Computing a tangent vector of a topological manifold thus requires: 1) definition of the manifold itself, 2) construction of set of charts covering the manifold, 3) definition of the basis functions *f*_*i*_, 4) definition of the scalar factors *α*_*i*_. We will discuss the final step in more detail below, but for now we note that steps 1-3 need only be carried out once per topological manifold. Once this has been done for a manifold, re-computing tangent vectors for different factors *α*_1_ simply requires carrying out simple calculations (which can be implemented in computer code) but no additional analytical work. Indeed, tangent vectors on any embedding of the topological manifold can also be effortlessly computed, as we will show next.

### Tangent vectors on the embedded manifold

The **embedding** of a topological manifold can intuitively be conceptualized as placing the manifold within another, higher dimensional, manifold or, like in this case, in a vector space, such that the resulting embedded manifold is still a topological manifold (i.e. without tears or self intersections). For a given sufficiently large embedding space, a manifold can be embedded in infinitely many ways, resulting in embedded manifolds with very different geometries (**Figure 1**A). Importantly, all embedded manifolds share the same topological structure as a result of being an embedding of the same topological manifold (e.g., any embedding of the plane is always a two dimensional surface in a higher dimensional space). Nevertheless, the widely different geometry of the embedded manifolds results in differently oriented tangent vectors, complicating the task of identifying tangent vectors for RNN design (**Figure 1**A).

This difficulty can be avoided simply by noting that given tangent vectors defined on a topological manifold and an embedding function *ϕ, ϕ* can be used to easily compute the corresponding tangent vectors on the embedded manifold. It is in fact possible to define a **pushforward** map *ϕ*^*\**^ : *T*_*p*_*M → T*_*ϕ*(*p*)_*ϕ*(*M*) assigning to each element *v*_*p*_ of the tangent vector space at a point on ℳ an element 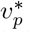 of the corresponding tangent vector space on the embedded manifold. An alternative approach, however, is to project the basis vectors of *TpM, e*_*i*_ = [*x*^−1^ ◦ *f*_*i*_] onto the embedded manifold 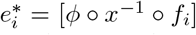 (**Figure 2**B). Then, 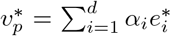 where the *α*_*i*_ are the same scalar factors as above.

The 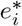 are valid tangent vectors according to the definition of tangent vectors as equivalence classes above. Creating manifold targeted RNNs, however, requires tangent vectors in the familiar form of *n*-dimensional vectors at a point to build a system of equations that can be solved to obtain the network’s connectivity matrix. These can be obtained by taking the derivative of the map *ϕ* ◦ *x*^−1^ ◦ *f*_*i*_ and evaluate it at *ϕ*(*p*) which can be done numerically since *ϕ x*^−1^ ◦ *f*_*i*_ is a parametrized curve in ℝ^*n*^(it is a map ℝ *to*ℝ^*n*^). The *n*-dimensional basis vectors 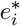 obtained through the derivative operation can be combined to give the *n*-dimensional vector form of 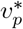 used for RNN fitting as described above.

Most manifolds of interest in neuroscience are low dimensional, and for these embedding maps onto ℝ^3^ are easy to define. For example, for the two dimensional manifold *S*^2^ (the sphere), *ϕ*(*p*) = *sin*(*p*_0_) *\* cos*(*p*_1_), *\* sin*(*p*_0_) *\* sin*(*p*_1_), *\* cos*(*p*_0_)) is an embedding (given an appropriate definition of the manifold’s set, see Methods) as it gives the familiar unit sphere centered at the origin. In general, however, embedding functions for arbitrarily large embedding spaces, like the ones of interest in neuroscience, are harder to define. To overcome this limitation, we note that an orthonormal (*n × k*) matrix *N* acts as an embedding of ℝ^*k*^ in ℝ^*n*^, for *k < n*. Thus, if an embedding *ϕ* of ℳ into ℝ^*k*^ exists, it is possible to embed ℳ in ℝ^*n*^as *Nϕ*(ℳ). This procedure can therefore be used to embed manifolds onto arbitrarily large state spaces, with the limitation that only embeddings that are possible in a lower dimensional space can be used, therefore reducing the range of geometries of the embedded manifold. Nevertheless, this procedure can produce rich and interesting embeddings of target manifolds in state space (see **Figure 2**C for embeddings in ℝ^64^). Importantly, the method for computing tangent vectors described here is agnostic to the way that embedding functions are obtained and can thus be used with embedding functions obtained through other methods.

### Tangent vector fields

In the preceding sections we demonstrated how a tangent vector can be defined as a linear combination of basis vectors of *T*_*p*_*M* or *T*_*ψ*(*p*)_*ψ*(*M*), which requires that *d*-many scalar coefficients *α*_*i*_ are specified. The choice of coefficients determines the orientation and magnitude of the tangent vector and thus the on-manifold dynamics of the RNN at that point. To obtain the desired dynamics then it is necessary to specify different coefficients for each point on the manifold. This can be achieved by defining a map *ψ* : ℳ → ℝ^*d*^ assigning a *d*-dimensional vector to each point on the manifold whose elements are the *α*_*i*_ coefficients. Thus *ψ* effectively assigns a tangent vector to each *p* ∈ ℳ, making it a **tangent vector field** and different choices of *ψ* define different vector fields on the same manifold (**Figure 2**D). Importantly, *ψ* is defined on the topological manifold such that the same vector field map can be used to generate RNNs with the same on-manifold dynamics but different embedded manifold geometry. On-manifold dynamics can therefore be defined on a *d*-dimensional space independently of the number of units in the network and how the *n*-dimensional dynamics unfold in state space (which depends, in part, on the choice of embedding function). Conceptually, *ψ* captures the latent variables dynamics solving a given task, which depends uniquely on the logical structure of the task itself (e.g., on number of latent variables and their dynamics; **Pollock and Jazayeri (2020); Jazayeri and Ostojic (2021); Maheswaranathan et al. (2019)**).

## Constructing targeted RNN

In this section we demonstrate that tangent vectors computed with the above procedure produce manifold targeted RNNs with the desired on-manifold dynamics. While **Pollock and Jazayeri (2020)** described a method for obtaining an RNNs connectivity matrix from a set of tangent vectors, here we describe a similar alternative procedure for achieving the same goal. Next, we use these procedure to create RNNs fitted to different manifolds and with different choices of embedding and vector field maps.

The RNNs used here are simple autonomous dynamical systems of *n* identical units whose dynamics are defined as:

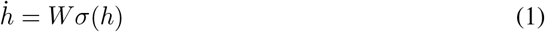

where 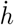 represents the first derivative of the network’s state *h* with respect to time, *W* is an *n × n* matrix whose entries define the strength of the connection between two units and *σ* is a non linear function (here *tanh*). Since the RNN dynamics are entirely specified by *W* once an initial condition is selected, the task of creating manifold targeted RNNs can be reduced to finding a *W* yielding the desired dynamics when the network’s state is initialized on the target manifold.

The network state corresponds to a point in the *n*-dimensional state space. For an RNN whose dynamics are confined to a target (embedded) manifold, this implies *h* = *ϕ*(*p*). Similarly, 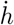 represents a velocity vector indicating how fast and in which direction the network state will evolve and must therefore be tangent to the target manifold at all times. Thus, 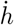 is a tangent vector of the embedded manifold 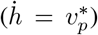 re-write We can then **equation 1** as:

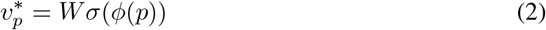

to reflect the notation described in the previous section.

Given a point on the topological manifold (and a vector field maps *ψ*) we can compute *ϕ*(*p*) and 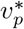 such that **equation 2** can be used to compute *W*. However, **equation 2** has *n* knowns and *n*^2^ unknowns. Thus, in practice, we sample *k*-many points on the topological manifold and build a system of equations:

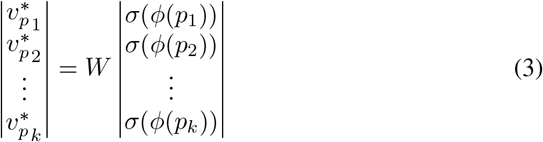

which can be used to solve for *W* using least squares.

This simple method can be used to construct RNNs whose dynamics lay on target manifolds in arbitrarily large state spaces(**Figure 3**). The manifold targeted RNNs thus obtained have low-rank connectivity matrices (with the rank matching the embedded manifold’s extrinsic dimensionality; **Figure 3**E), yet they show minimal off-manifold deviations over time (**Figure 3**B) and can display rich and accurate on-manifold dynamics (**Figure 3**C,D). Indeed the approach outlined here can be used to easily construct RNNs with task-solving dynamics **Figure 3**F. The appropriate choice of on-manifold dynamics allows the network to solve a 2-bit memory task by using four fixed points attractors to store its internal representation of the inputs. New inputs can update this representation by moving the network state between different attractors. Furthermore, the dynamics are stable with respect to repeated, redundant, inputs which do not push the state beyond the current attractor’s basin of attraction. While gradient-descent algorithms are used to train networks to perform this kind of task **(Sussillo and Barak, 2013)**, here careful choice of the dynamics manifold and on-manifold dynamics yields a task-solving network directly.

**Figure 3.**
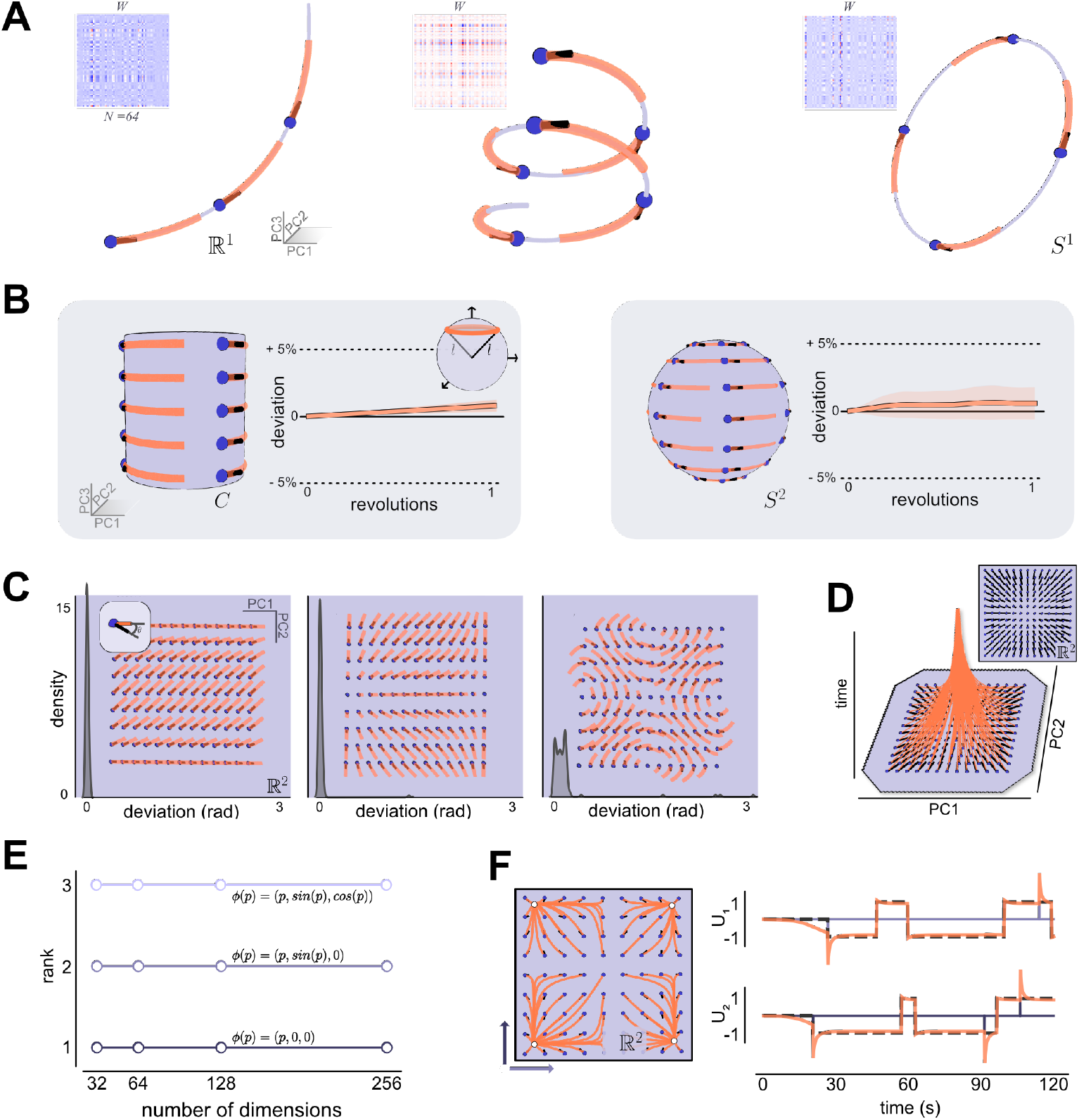
**A**. RNN dynamics (salmon traces) for RNNs targeted to one dimensional manifolds embedded in ℝ^64^ visualized in PCA space (see Methods). Different traces show the RNN dynamics when the RNN was initialized to a different initial condition (blue dots). Insets show the connectivity matrix of RNNs producing the dynamics shown in figure. **B**. Quantification of dynamics drift for the cylinder (left) and sphere (right) manifolds in ℝ^64^. RNNs were created with dynamics as shown on the left of each panel such that the distance from the origin of state space would remain constant over time (see inset). The plots on the right on each panel show the change in distance from the origin over one complete revolution across different starting points. **C**. Quantification of dynamics accuracy for different vector fields on the plane manifold in ℝ^64^. Three different vector fields were defined for the same manifold, producing different RNN dynamics (shown in the background of each panel). The angle between the RNN dynamics and the tangent vector at the point in which the RNN was initialize was quantified (see inset) and shown as kernel density estimate distributions. **D**. Dynamics fixed points. Visualization of the dynamics of an RNN fitted using a vector field which specified a single attractor point (see inset). The plot shows the dynamics evolving in the plane spanned by the first two principal components (see Methods) over time. **E**. Connectivity matrix rank. The rank of the connectivity matrix *W* of networks of different dimensions fitted to three different embeddings of the line manifold in ℝ^*n*^. **F**. Task solving dynamics. Left, vector field and RNN dynamics on ℝ^2^. Arrows indicate the two inputs vectors *u*_1_ and *u*_2_. Right, example trial. Purple lines show the two inputs, dotted lines the expected network outputs. Solid orange lines show the actual network outputs.

## Discussion

In this manuscript we have leveraged topology and differential geometry to simplify the task of computing vectors tangent to a manifold in state space. These vectors can be used with local-geometry based RNN engineering algorithms **(Pollock and Jazayeri, 2020)** to create RNNs with dynamics that unfold along the target manifold and have rich, task-solving, on-manifold dynamics. Our approach facilitates the computation of tangent vectors by carrying it out on a topological manifold prior to embedding it in state space. This has the advantage that once a tangent vector is defined on the topological manifold, the corresponding vector can be computed on any embedding of the manifold in state space.

By using the language of differential geometry we can separately define the geometry of the embedded manifold (which depends on the embedding function) and the on-manifold dynamics (defined by a vector field over the topological manifold), as well as the manifold topology itself. Conceptually, the topology of a network’s activity manifold and its on-manifold dynamics are determined by the logical structure of the task being performed, and are shared across networks with different architectures **(Maheswaranathan et al., 2019)**. On the other hand, the geometry of the embedded manifolds is constrained by the properties of the individual network (i.e., choice of hyperparameters) and largely independent of the computation being performed. For example, networks with different non-linear activation functions (e.g.: *tahn* vs *ReLu*) might require that the embedded manifold is confined to different sub-regions of state space. Differential geometry provides a language for describing manifold topology and geometry independently, thus mirroring the independent nature of these phenomena.

The intrinsic topology of neural manifold and on-manifold dynamics are increasingly regarded as crucial for understanding neural computations **(Jazayeri and Ostojic, 2021; Darshan and Rivkind, 2021; Chung and Abbott, 2021)**. Having a conceptual framework for exploring these ideas is thus crucial for gaining further insights into the link between network connectivity, dynamics and computation. Given the intrinsically geometric nature of network dynamics, we believe that differential geometry should be a fundamental element of such framework. It allows for deep and precise understanding of several key concepts in neural dynamics (e.g. manifold topology vs embedded geometry), thereby providing a promising venue for the abstraction of fundamental mechanistic explanations of computation across neural networks with different properties. The approach presented here is a step in applying topology and differential geometry based approaches to investigate the link between connectivity, dynamics and computation, as well as to enable the application of methods such as EMPJ **(Pollock and Jazayeri, 2020)** to a wider class of problems in neuroscience and machine learning.

## Methods

### Code availability

All work presented in this manuscript was carried out using custom python code. The code, included scripts to replicate all figures, is available at the GitHub repository: https://github.com/FedeClaudi/manyfolds. The python code makes use of open source science software python packages including numpy, scikit and matplotlib **(Harris et al., 2020; Pedregosa et al., 2011; Hunter, 2007)**.

### Manifolds

#### manifolds definitions

Topological manifolds are defined by a choice of set *M* and topology. Here we assume the standard topology throughout, the following sets were used to define each manifold:

- **line** ℝ^1^: *M* = [0, 1]
- **circle** *S*^1^: *M* = [, 2*π*]
- **plane** ℝ^2^ : *M* = [0, 1] × [0, 1]
- **cylinder** *C*: *M* = [0, 2*π*] × [0, 1]
- **sphere** *S*^2^: *M* = [0, *π*] × [, 2*π*]

where × denotes the Cartesian product.

#### manifold embeddings

All manifolds were embedded in ℝ^64^except for **Figure 3**E. The embedding was achieved in two steps as described in the text: a function *ϕ* : ℳ → ℝ^3^ was used to embed the manifolds in ℝ^3^ and a random orthonormal 3 × 64 matrix acted as an embedding map ℝ^3^ *→* ℝ^64^.

The following embedding functions *ϕ* were used:

- ℝ^1^ as **helix**.

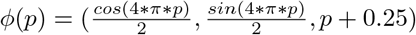
- ℝ^1^ as **line**. *ϕ*(*p*) = (*sin*(2*p*)−0.5, 2*sin*(*p*)− 1, -4*cos*(*p*) + 3)
- *S*^1^ as curved **circle**.

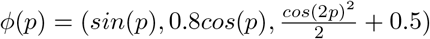
- *S*^1^ as bent **circle**.

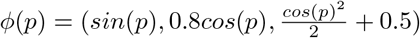
- *C* as **cone**.

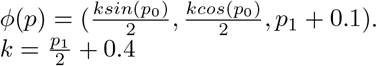
- *C* as **cylinder**.

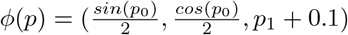
- ℝ^2^ as curved **plane**. *ϕ*(*p*) = 2(*p*_0_, *sin*(*p*_1_), 0.4(*p*_1_ − *p*_0_)^2^)
- ℝ^2^ as flat **plane**.

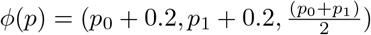
- *S*^2^ as unit **sphere**. *ϕ*(*p*) = (*sin*(*p*_0_)*cos*(*p*_1_), *sin*(*p*_0_)*sin*(*p*_1_), *cos*(*p*_0_))

#### visualizing embedded manifolds

Three dimensional visualizations of embedded manifolds and RNN dynamics were realized with the python package vedo **(Musy et al., 2021)**. A set of *m* (25-1000) points were sampled from the topological manifold and the coordinates in embedding space were computed. Then a PCA model model was fitted using the scikit package **(Pedregosa et al., 2011)** and the first three principal components was used to reduce the dimensionality of the point cloud to three dimensions. The manifold’s surface was reconstructed in PCA space using algorithms implemented in vedo. For **Figure 3**D, the first two principal components were used and the third dimension represented time.

### Manifold charts ad basis functions

To define a set of charts covering the topological manifold one or two charts per manifold were used. When two charts were used, the chart set *U*_*i*_ and map *x*_*i*_ were carefully selected such that *x*_*i*_(*U*_*i*_) was homeomorphic to the same set in ℝ^*d*^for each chart, simplifying the definition of basis functions. The basis functions were simply defined as maps *f*_*i*_ : [0, 1] *→ x*_*i*_(*U*_*i*_) parallel to one axis of the local coordinate space. For instance for a two dimensional manifold with *x*_*i*_(*U*_*i*_) ≅ [0, 1] × [0, 1], the first basis function for a point *p* ∈ ℳ is defined as *f*_1_(*λ*) = (*λ, x*(*p*)).

The following charts and basis functions were used:

- For ℝ^1^ a single chart (*x, U*) was defined with *U* = *M* and *x* := *x*(*p*) → 2*p*.
- for *S*^1^, two charts were used with *U*_1_ = [0, *π*], *U*_2_ = [*π, 2π*] and *x*_1_ = *id, x*_2_ := *x* (*p*) = *p* – *π*
- for ℝ^2^ a single chart was used with *U* = *M* and *x* = *id*.
- for *S*^1^, two charts were used with *U*_1_ = [0, *π*] × [0, *π*], *U*_2_ = [0, *π*] × [*π, 2π*] and 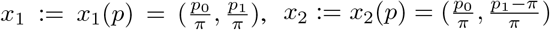.
- for *C* two charts were used with *U*_1_ = [0, *π*] × [0, 1], *U*_2_ = [*π*, 2*π*] × [0, 1] and 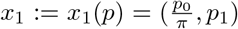 and 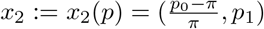.

### Vector fields

Tangent vector fields maps (*ψ*) were used to specify the magnitude and orientation of tangent vectors. For **Figure 2**D, the following vector field maps were used:

- 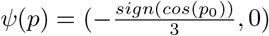
- 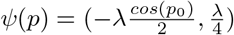 with *λ* = 1.5 − *abs*(*cos*(*p*_0_))
- 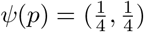

For **Figure 3**A, the vector field was *ψ*(*p*) = 1 while in panel **B** the vector field used was *ψ*(*p*) = (0, 1) for the sphere and *ψ*(*p*) = (1, 0) for the cylinder. In panel **C** the three vector fields were:

- 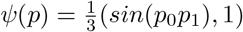
- 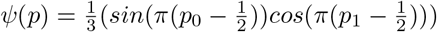
- 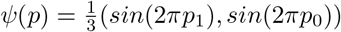

For panel 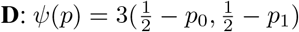 The vector field for panel **F** is described in the section *Task solving dynamics* of the methods.

### Tangent vectors computation

To compute the tangent vector at a point *ϕ*(*p*) on the embedded manifold, the position *x*_*i*_(*p*) of the point in the chart representation was computed (given a chart (*x*_*i*_, *U*_*i*_) such that *p ∈ U*_*i*_). Next, the basis functions were defined as described above to obtain a one dimensional curve in *x*_*i*_(*U*_*i*_) ⊂ ℝ^*d*^ for each basis. Next, the projection of the basis functions onto the embedded manifold was computed as *ϕ ◦ x*^−1^ ◦ *f*_*i*_ yielding one dimensional curves in ℝ^*n*^. The derivative of these curves with respect to the parameter *λ* of *f*_*i*_ was computed and evaluated at the point *ϕ*(*p*), to obtain the basis tangent vector. Finally, the tangent vector field map *ψ* was evaluated at *p* to obtain the coefficients for the linear combination of basis vectors and thus compute the tangent vector.

### RNN creation

The procedure for generating manifold targeted RNNs is described in the main text. Here we note that for each manifold a set of *k* = 3 − 100 equally spaced points was used to build the system of equations whose solution, obtained via least squares using numpy, gave the RNN connectivity matrix *W*,

### Dynamics drift

To estimate how much the RNN dynamics drifted off the surface of the target embedded manifolds over time, 10 RNNs were fitted to the cylinder and sphere manifolds each using the embedding and vector field maps as described above. Each RNN was initialized at 25 different locations on the embedded manifold and allowed to evolve until it completed an entire revolution around the manifold and returned to its original position. The average and standard deviation of the normalized distance from the origin of state space for all RNNs and all starting positions on a given manifold was computed and shown in figure.

### Dynamics accuracy

To estimate how accurately RNN dynamics matched the direction prescribed by tangent vector fields, three different vector fields on the flat plane embedded manifold were used, as described above. For each, 10 RNNs were fitted to it and initialized at 81 different locations on the manifold from where they were allowed to evolve for the simulation time equivalent of five seconds. The angle between the vector from the initial location and the final RNN state and the tangent vector at the initial location was computed for each initialization.

### RNN matrix rank

To estimate the rank of manifold targeted RNNs’ connectivity matrices (**Figure 3**W), RNNs with different number of units (n=32, 64, 128 and 256) were fitted to the line manifold ℝ^1^ in ℝ^*n*^with one of the following embedding maps:

- *ϕ*_1_(*p*) = (*p*, 0, 0), a straight line.
- *ϕ*_1_(*p*) = (*p, sin*(*p*), 0), a planar curve.
- *ϕ*_1_(*p*) = (*p, sin*(*p*), *cos*(*p*)), a three dimensional curve.

to produce embedded manifolds with different extrinsic dimensionalities. The rank of the connectivity matrix *W* of each RNN was then estimated in python, using the numpy library.

### Task solving dynamics

#### 2-bit memory task design

The 2-bit memory task was created by adapting the 3-bit memory task used in other works **(Sussillo and Barak, 2013; Maheswaranathan et al., 2019)**. In brief, two independent and time-varying inputs were produced. At each time step the inputs could assume values of 1, -1 or 0. The task’s goal consists in keeping track the last non-zero value received from each input. Given two inputs and two possible values for each, four possible combinations are possible. The network must keep an internal representation of the current combination and updated it correctly in response to new non-zero inputs. This include ignoring redundant inputs (e.g., two consecutive 1s) which should not change the network’s internal representation.

#### RNN design

As the starting point for designign the task-solving RNN dynamic, we used the solution discovered by training networks on the 3-bit memory task **(Sussillo and Barak, 2013; Maheswaranathan et al., 2019)**. The presence of two latent variables (the ‘state’ of each input) naturally suggests the use of a two dimensional manifold, the absence of a periodic structure in the inputs suggests that a plane would be ideal. For simplicity, we embedded ℝ^2^ in ℝ^*n*^(n=64) as a flat plane by: 1) selecting a random *n*-dimensional vector (*v*_0_), 2) selecting a second random vector orthogonal to the first (*v*_1_) and 3) using the embedding function 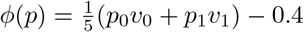.

The requirement that the network be able to maintain four stable states in the absence of stimuli suggests that four fixed point attractors should be included in the on-manifold dynamics. Stimuli should push the dynamics from one attractor state to another to change the network’s internal memory, but repeated stimuli should not push the network state outside the current attractor’s basin of attraction. To achieve this the following tangent vector field was used: *ψ*(*p*) = 0.6(−*sin*(2.5*π*(*p*_0_ − 0.1)), −*sin*(2.5*π*(*p*_1_ − 0.1)))

The RNN dynamics **equation 1** does not include external inputs. To create a task-solving RNN we modified **equation 1** to:

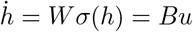

where *u* is the two-dimensional inputs vector and *B* is the (*n* × 2) input matrix describing how each input affects each unit in the network. We defined

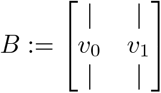

such that each input moved the network along one of the ‘sides’ of the plane manifold.

The network output was a two dimensional vecotr defined as *y* = *B*^−1^*h* thus translating the position along each ‘side’ if the embedded plane into a scalar output.

## Mathematical Appendix

### Topological manifold and topology

A **topological manifold** ℳ is an Hausdorff connected topological space locally homeomorphic to ℝ^*d*^. Thus, for every point *p ∈ ℳ*, there exists a neighborhood *U* with *p* ∈ *U* such that *U* is homeomorphic to ℝ^*d*^with *d* constant for every point in the manifold. Then *d* also specifies the dimensionality of the manifold.

A **topological space** is defined by a set of points *M* and a **topology** 𝒪. The set is generally an interval *I*⊂ ℝ or the Cartesian product of *d*-many such intervals. Different sets can be used to define the same manifold, for instance the line ℝ^1^ can have *M* = [0, 1] or *M* ′ = [1, 2] as underlying sets. The topology 𝒪 is defined as a subset of the powerset of *M* : (*P*)(*M*) such that: 1) Ø, *M*, ∈ 𝒪 2) unions of elements 𝒪 of are also in and 3) intersections of elements of 𝒪 are also in 𝒪.

The **standard topology** is intuitively conceptualized as the topology of Euclidean space ℝ^*d*^. More precisely, for every *x ∈ M*, and *rin*ℝ^+^, define the open ball of radius *r* centered at *x* as the set 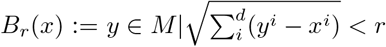. Then, an open subset *U* is in the standard topology of ℝ^*d*^if there’s a ball of some, possibly small, radius contained in *U*.

### Tangent vector

Given a parametrized curve *γ* : ℝ ⊃ *I* → ℳ and a point *p* ∈ ℳ, then two equivalent definitions of **tangent vectors** can be given. A tangent vector can be defined as the directional derivative operator at *p* along *γ* or, alternatively, as the equivalence class of parametrized curves through *p* sharing the same directional derivative. The two definitions are equivalent, there’s a one-to-one relationship between the directional derivative operators and the equivalence classes, in this work we used the latter definition.

### Tangent vector space

The tangent vectors at a point *p* ∈ ℳ define a (tangent) vector space *T*_*p*_*M* with well-defined operations of vector addition and scalar multiplication (technically *T*_*p*_*M* is an algebra since an operation of vector multiplication is also defined). A fundamental result in differential geometry proves that *dim*(*M*) = *dim*(*T*_*p*_*M*), that is *T*_*p*_*M* and ℝ^*d*^ are isomorphic as vector spaces. Given a chart (*x, U*) with *p ∈ U*, the basis of *T*_*p*_*M* can be constructed by defining a set of *d*-many curves *f*_*i*_ in the chart representation of *U*, and projecting them onto the manifold using (^−1^*x*). These curves then act as representative for the equivalence classes that are the basis vectors. The basis functions are maps *f*_*i*_ : ℝ ⊃ *I* → *x*(*U*) defined to be parallel to the *i*^*th*^ axis of the local coordinates frame determined by the char.

### Chart

A chart of a *d*-dimensional manifold ℳ with set *M* is a pair (*x, U*) with *U* ∈ 𝒫(*M*) and *x* : *U → x*(*U*) ⊂ ℝ^*d*^ a bijective homemorphism of *U* onto a subset of ℝ^*d*^. The existence of *x* is ensured by the definition of a topological manifold as locally homeomorphic to ℝ^*d*^. The component functions *x*^*i*^ : *U* → ℝ are called the coordinates of *p* with respect to the chart (*x, U*) and the chart establishes a local coordinates system around *p*. One or more, potentially overlapping, charts can be defined such that the union of the charts’ set covers *M*.

### Embedding

An **embedding** is a map between topological manifolds *ϕ* : ℳ → 𝒩 that is an **immersion** and such that the mapped manifold *ϕ*(ℳ) is a submanifold of the target manifold (i.e., it’s a manifold *L* whose set ℒ is a subset of the set *N* of 𝒩). An immersion is a smooth map *ϕ* between manifolds such that its derivative (the **pushforward** map *ϕ*^*\**^) is injective at every point *p* ∈ ℳ In this work, the target manifold is the vector space ℝ^*n*^which is a topological manifold with additional operations of vector addition and scalar multiplication defined.

### Pushforward

Given an map between manifolds *ϕ* : ℳ → 𝒩, then the **pushforward** (or derivative) of *ϕ, ϕ*_*\**_ is a map between tangent vector spaces of the source manifolds to tangent vector spaces on the target manifold: *ϕ*_*\**_ : *T*_*p*_*M → T*_*ϕ*_*N*. Given the definition of tangent vectors as equivalence classes of curves, the pushfoward of the vector [*γ*], *ϕ*_*\**_*γ*] is [*ϕ* ◦ *γ*]_*ϕ*(*p*)_.

### Tangent vector field

Given a manifold ℳ, the manifold together with all the vector spaces *T*_*p*_ℳ at each point in ℳ form a bundle of topological manifolds: *TM* : = ∪ _*p*∈*M*_ *T*_*p*_*M*. A vector field, or tangent vector field, is a section *ψ* of the topological bundle. That is *ϕ* is a map *ψ* : ℳ → *T*_*p*_*M* assigning to each point *p* on the manifold a tangent vector (an element of the tangent vector space *T*_*p*_*M*).

